# Efficient Search for Circuit Structure by ‘Smooth-Index’ Matrix-Reordering

**DOI:** 10.1101/2023.09.26.559501

**Authors:** Alexander Borst, Winfried Denk

## Abstract

Volume electron microscopy together with computer-based image analysis are yielding neural circuit diagrams of ever larger regions of the brain [1-10]. These datasets are usually represented in a cell-to-cell connectivity matrix and contain important information about prevalent circuit motifs allowing to directly test various theories on the computation in that brain structure [11,12]. Of particular interest are the detection of cell assemblies and the quantification of feedback, which can profoundly change circuit properties. While the ordering of cells along the rows and columns doesn’t change the connectivity, it can make special connectivity patterns recognizable. For example, ordering the cells along the flow of information, feedback and feedforward connections are segregated above and below the main matrix diagonal, respectively. Different algorithms are used to renumber matrices such as to minimize a given cost function, but either their performance becomes unsatisfying at a given size of the circuit or the CPU time needed to compute them scales in an unfavorable way with increasing number of neurons [13-15]. Based on previous ideas [16-18], we describe an algorithm which is effective in matrix reordering with respect to both its performance as well as to its scaling in computing time. Rather than trying to reorder the matrix in discrete steps, the algorithm transiently assigns a real-valued parameter to each cell describing its location on a continuous axis (‘smooth-index’) and finds the parameter set that minimizes the cost. We find that the smooth-index algorithm outperforms all algorithms we compared it to, including those based on topological sorting.

**Author Summary:** Connectomic data provide researchers with neural circuit diagrams of ever larger regions of the brain. These datasets are usually represented in a cell-to-cell connectivity matrix and contain important information about prevalent circuit motifs. Such motifs, however, only become visible if the connectivity matrix is reordered appropriately. For example, ordering the cells along the flow of information, feedback and feedforward connections are segregated above and below the main matrix diagonal, respectively. While most previous approaches rely on topological sorting, our method treats the discrete vertex indices as real numbers (‘smooth-index’) along independent parameter axes and defines a differentiable cost function, thus, allowing gradient-based algorithms to find a minimum. The parameter set at this minimum is then re-discretized to reorder the connectivity matrix accordingly. We find our method to scale favorably with the circuit size and to outperform all algorithms we compared it to.

## Introduction

A natural way to order the elements of a circuit is along the direction of information flow. This way, import features of the circuit become immediately visible in the respective connectivity matrix: synfire chains will form diagonals of the matrix, and squared blocks along the diagonal will indicate the presence of cell assemblies. Arranging the outputs of each neuron within the columns of the connectivity matrix, the ordering will put forward synapses in the lower and recurrent connections in the upper triangle. Isolating recurrent synapses can be achieved by ordering the connectivity matrix such as to push as many entries to the lower triangle as possible, a so-called ‘minimum feedback arc set’. We focus on recurrency as a specific property of neural circuits because of the importance recurrent synapses have for the temporal processing properties of the circuit. In networks without feed-back, the eigenvalues of the corresponding dynamical system matrix *A* = *T*^−1^(*M* − *I*) - with T being the matrix holding the cellular time-constants along the diagonal, M being the connectivity matrix and I the identity matrix - are the negative inverse of the cellular time-constants [14]. In contrast, in networks with feed-back, the eigenvalues depend not only on the cellular time-constants, but in addition on the connectivity parameters [14,19-21]. Thus, feed-back generally generates new time constants and thereby allows for signal processing at time scales that can go beyond the ones of the isolated elements by orders of magnitude, thereby providing a substrate for working memory [22-24], path integration [25,26] or delay lines in the context of motion vision [27,28]. Within a given connectomic data set, however, neurons usually are not numbered according to the main flow of information within the circuit. In order to retrieve information from the dataset, neurons must therefore be reordered with a specific objective in mind. Reordering of a square matrix with size N x N starts with an index list π = (π_1_, π_2_ … π_*N*_) which assigns each circuit element i the new index π_*i*_. From this, a permutation matrix *P* is created by permuting the columns of the identity matrix accordingly, and the reordered connectivity matrix *R* is obtained by *R* = *PMP*^−1^. Since renumbering preserves the connectivity information, the graphs resulting from renumbering are all isomorphic. Different algorithms exist to reorder matrices [13-15]. As a fundamental common property, they all retain the discrete nature of the vertex indices. This way, any property defined to be optimized is not a differentiable function along any parameter axes, but rather a discrete value depending on a given permutation. Since a full enumeration of all permutations grows with O(N!), a brute force approach fails quickly as N increases. The problem presents itself as a sorting problem where each of the different algorithms apply a different, particular strategy.

## Results

Our approach is different from topological searches but rather based on graph layout techniques [16-18] (Fig.1A). We first release each vertex index from being a discrete integer number and turn it into a smooth real number z_i_ that indicates the respective vertex position on a continuous axis. Next, each vertex position is considered to be a parameter value along an independent axis. Thus, the cost to be minimized becomes a function of an N-dimensional parameter space. Although the number of possible solutions, i.e. all permutations of N, represent only a small subspace of the new search space N^N^, this transition brings the advantage that the cost function can become differentiable, allowing gradient descent methods to be applied to search for a minimum. A recurrent connection is characterized by the fact that the position z_m_ of a presynaptic neuron m is larger than the position z_n_ of a postsynaptic neuron n. However, the number of recurrent connections varies in a discrete step at the point where z_m_ = z_n_. Hence, it is not a differentiable function of z. We therefore chose the average length z_m_ – z_n_ of all recurrent connections as a proxy for the number of recurrent connections, and, by taking a saturating function of the length, such as the logistic function, reduced the stronger influence of longer connections over shorter ones (Eq.1, Methods). To restrict our solutions to the permutation subspace, we require, as an additional criterion to the original cost function, each position to be different from all other positions. To this end, we define an additional ‘Pauli term’ as the mean of the squared differences between the positions and their rank, i.e. their place within the sorted array of positions (Eq.5, Methods). To return to discrete vertex indices, as the final step, the permutation list π that reorders the original matrix is obtained as the vector containing the arguments of the rank-sorted parameters at the minimum of the cost function.

**Fig 1.**
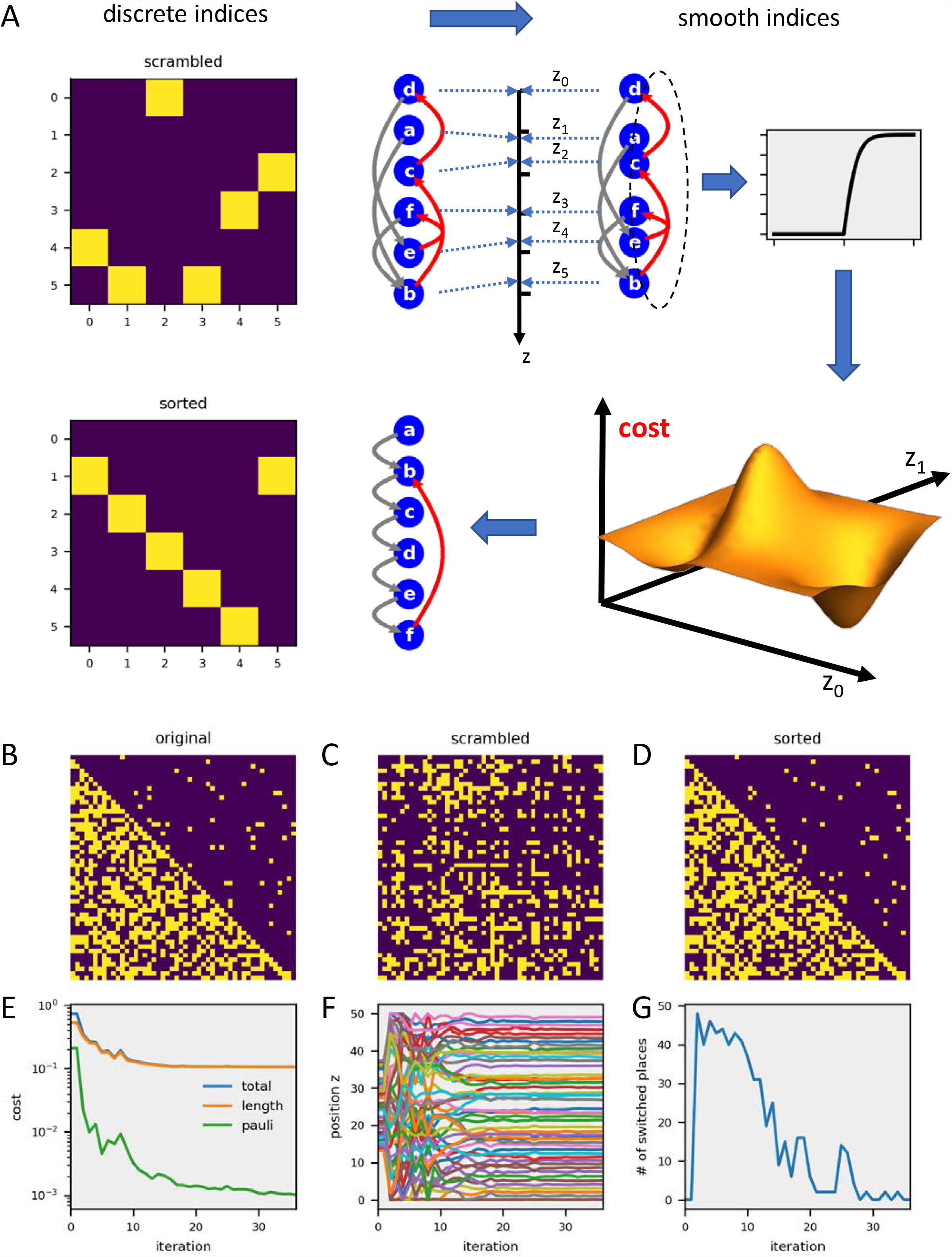
Working scheme and example performance of the smooth-index algorithm. **A** Starting from a ‘scrambled’ circuit, the algorithm treats the indices as smooth values along independent parameter axes and minimizes a given cost function, in this case all recurrent connections. **B** Original matrix with a few recurrent connections. **C** Scrambled matrix, obtained by random index permutation. **D** Matrix resulting from smooth-index sorting. **E** Different cost functions as the algorithm performs the gradient descent. **F** Positions of all vertices z_i_ during gradient descent. **G** Number of switches, calculated as the number of differences between two consecutive rank-sorted position vectors, during gradient descent.

We implemented the smooth-index algorithm in Python using the Scipy minimize function [29]. For a time-efficient search, we calculated the gradient or ‘Jacobian’ of the cost function (Eq.2 & Eq.6, Methods), which holds all first-order partial derivates of the cost function along each parameter z_n_. The cost function as well as the Jacobian were passed on to the Scipy minimize function with randomly initialized parameter values z. As shown in Fig.1B-G for a circuit with 50 neurons, the algorithm works well and isolates a similar number of recurrent synapses as is found in the original connectivity matrix. To test the efficiency of the sorting algorithm quantitatively, we applied it to matrices of different sizes, from 10 up to 10 000 neurons. Each circuit was constructed randomly with a given probability for entries in the lower and upper triangle. To quantify the performance, we calculated the number of non-zero entries in the upper triangle of the original connectivity matrix, i.e. before scrambling, relative to the number after smooth-index sorting. The results demonstrate that the algorithm reorders the scrambled connectivity matrices with a high degree of fidelity, in particular for large numbers of neurons (Fig.2A, blue line). Topological sorting according to the out-degree of the nodes performs significantly worse (Fig.2A, red line). For the smooth-index algorithm, the CPU time needed to compute the reordered matrix on a standard desktop computer grows from about 0.1 seconds for N=10 to about 1000 seconds for N=10 000. For larger networks, the CPU time roughly scales with O(N^2^) (Fig.2B, blue line). Not astonishingly, out-degree sorting is always faster (Fig.2B, red line).

**Fig 2.**
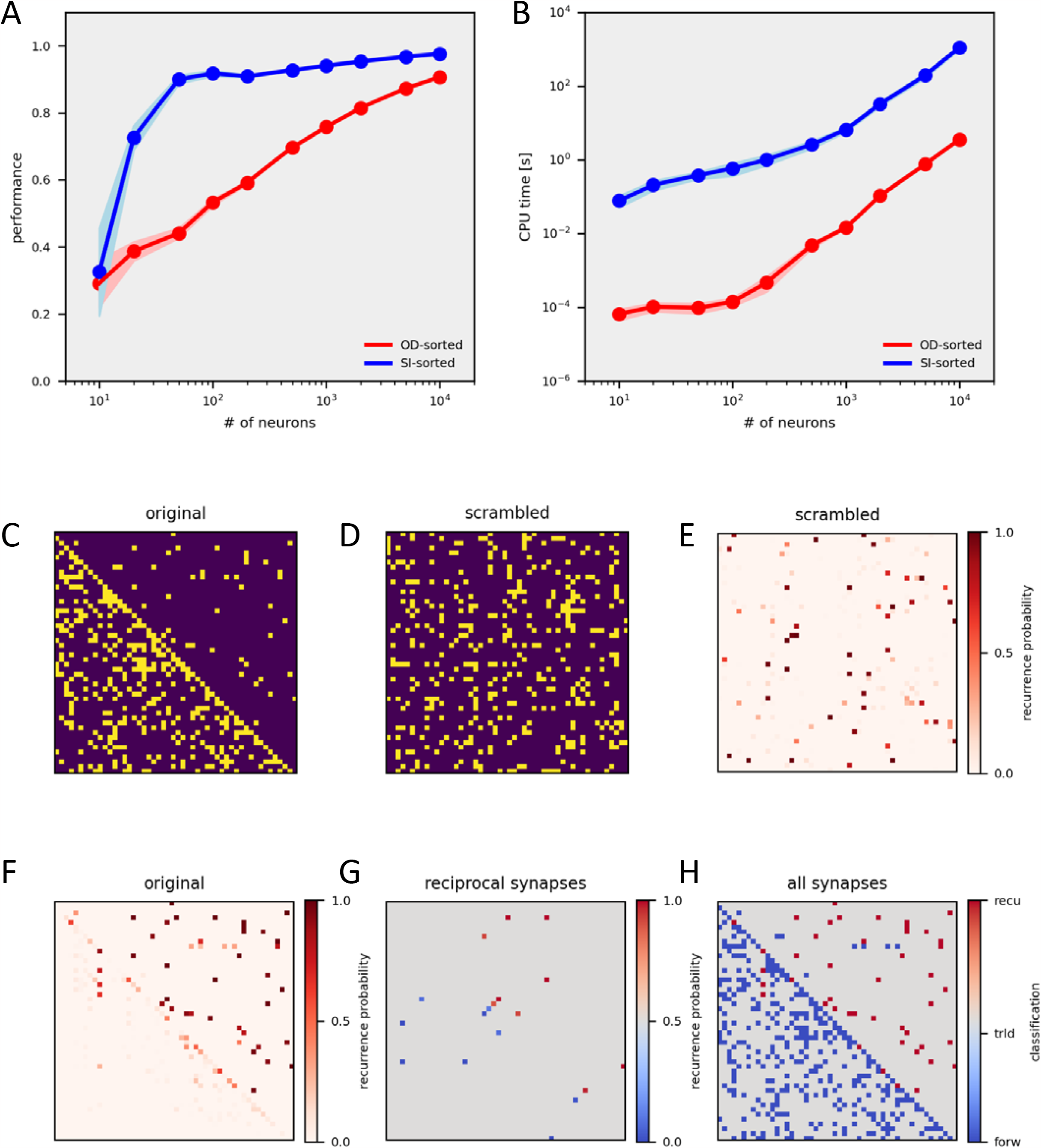
Performance of the smooth-index algorithm. **A** Performance of the algorithm as a function of the number of neurons. Plotted is the relative number of non-zero entries in the upper triangle of the original connectivity matrix with respect to the one after smooth-index sorting. **B** CPU time needed for out-degree sorting (in red) and for smooth-index sorting (in blue) as a function of the number of neurons. Data in A and B represent the mean +-standard deviation (shaded area) obtained from 10 sorting runs. **C-H** Identification of recurrent synapses by the smooth-index algorithm. **C** Original matrix. **D** Scrambled matrix. **E** Matrix holding the probability of each connection as being classified as recurrent from 1000 sorting runs. **F** Same as E, but after reversed scrambling. **G** Recurrence probability of all reciprocal connections. **H** Original connectivity matrix (in blue) with identified recurrent connections in red (probability threshold = 0.45).

However, yielding a similar number of recurrent connections does not mean that these connections are identical to the ones in the original, unscrambled matrix. In order to see whether we can also identify recurrent synapses, we constructed a single connectivity matrix with about 12% recurrent synapses (Fig.2C), scrambled it once (Fig.2D) and subjected it to the smooth-index sorting 1000 times. With each run being randomly initialized, different minima were likely do be found each time. From all these different runs, we constructed a matrix which holds the probability values by which each connection was classified as recurrent (Fig.2E). Reversing the scrambling revealed that most recurrent synapses of the original matrix were classified as recurrent with a high probability (Fig.2F). Here, an especially interesting case are reciprocal connections between two cells. Given the fact that if one of the connections is feed-forward, the other must be recurrent, one might naively expect that, for reasons of symmetry, each of these reciprocal connections is classified as recurrent with a probability of 0.5. However, contrary to this expectation, looking at all pairs of reciprocal synapses, only one of the connections had a high, while the other had a low probability of being classified as recurrent (Fig.2G). Obviously, taking into account the full connectivity, the symmetry between reciprocally connected neurons breaks. In the end, our method correctly identified 82% of all recurrent connections of the original matrix (Fig.2H). We conclude that our method allows us to not only calculate the degree of recurrency, i.e. how many recurrent connections exist in a given network, but also to successfully identify these recurrent synapses and isolate them from the feed-forward ones.

We finally explored whether the feedback detection problem is unique in being susceptible to smooth-index optimization. Another reordering problem with relevance to Neuroscience is the unmasking of clusters and processing chains. In order to apply the smooth-indexing method to identify properties like a block-diagonal structure or a limited band-width, we define the cost function as the mean squared length of all connections, irrespective of their sign, similar to an earlier approach [18] (Eq.3, Methods). Its gradient is given by Eq.4 (Methods). The algorithm successfully identifies the original structure of a band-width limited connectivity matrix (Fig.3A,B) as well as of a block-diagonal connectivity matrix (Fig.3C,D) with a high fidelity and outperforms the standard reverse Cuthill-McKee algorithm [30] over circuit sizes up to 10 000 neurons.

**Fig 3.**
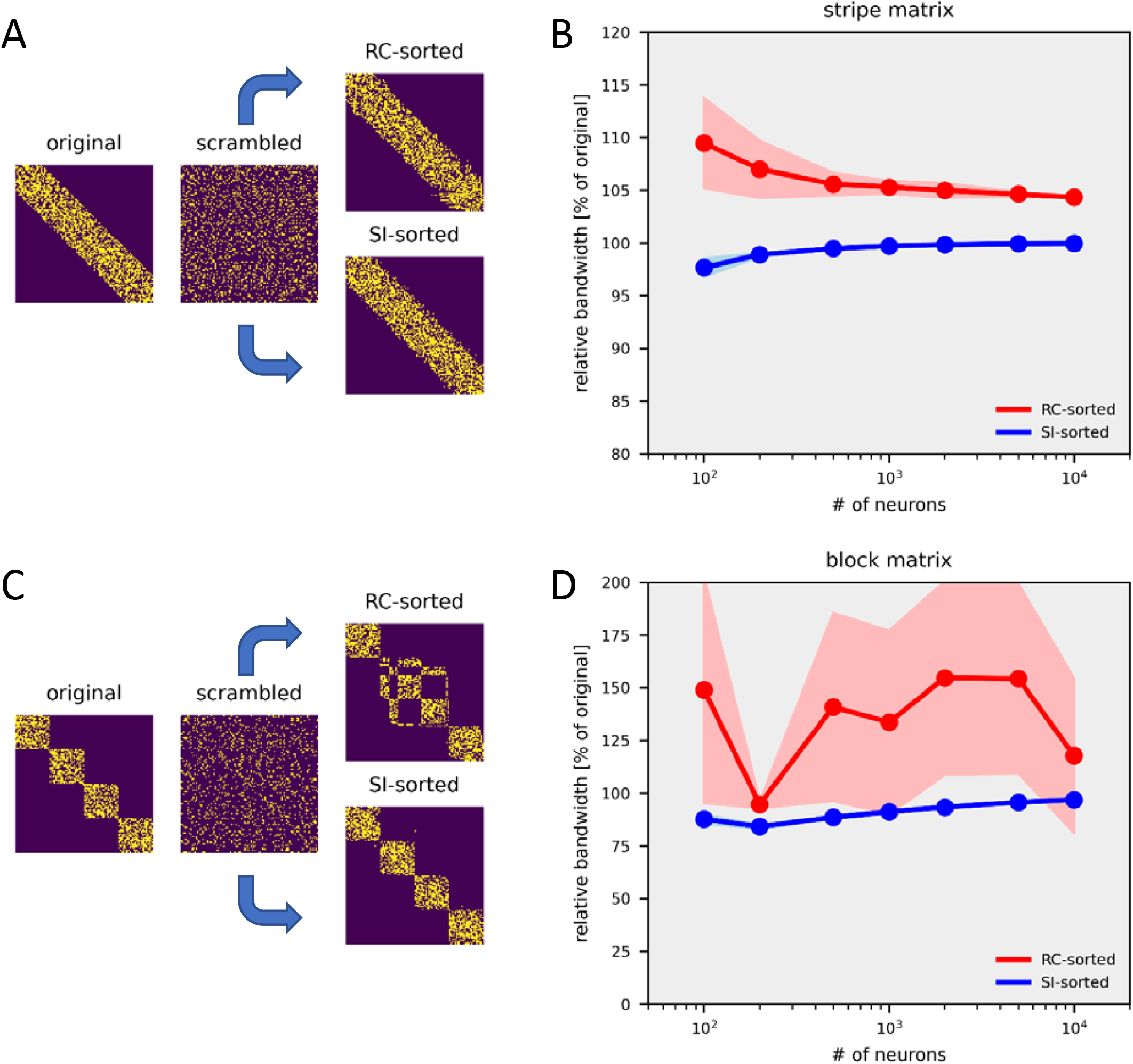
Application of the smooth-index to bandwidth-limited matrices. (**A**,**B**) *and to block-diagonal matrices* (**C**,**D**). In A and C, one example sorting is shown for a 50 x 50 matrix. In B and D, the performance of both algorithms is quantified as the bandwidth of the reordered matrix relative to the original matrix. The performance is calculated as the average obtained from 10 runs +-the standard deviation (shaded area).

## Discussion

Searching for an optimum as a function of permutations within a given range is a hard problem because the number of possible permutations grows with N-factorial. Various approaches have been taken in the past to deal with this problem like the Cuthill-McKee algorithm [30], page-rank algorithms [31] or flow-graph approaches [32,33] (for review, see [13-15]). Depending on the particular objective and the specific algorithm, they are based on reordering the graph nodes according to their degree, i.e. the number of edges each node has, or partitioning the original graph into subgraphs according to similarity of rows and columns in the respective connectivity matrix. At the end, in one way or the other, these algorithms all test random permutations within the subgraphs in an iterative way, always assessing the reordering according to a given quality criterion. As a common property, all these algorithms keep the vertex indices as discrete values. Seung [18] was first to apply graph layout techniques to reorder matrices. Defining a quadratic cost function for bandwidth minimization, he derived the gradient, set it to 0 and solved the linear matrix equation by using the Moore-Penrose-inversion. While this approach is fast and works reasonably well to identify connection chains and block-diagonal structures, it is, however, not applicable to other, non-quadratic cost functions, like the one needed to isolate recurrent connections. Here, our approach described above is more general and allows for bandwidth minimization as well as for isolation of recurrent synapses, with only a small excess of recurrent connections compared to the original circuit. While all our examples above used simple graphs with corresponding binary connectivity matrices, smooth-index sorting can easily be extended to weighted graphs, preserving the strength of connections. In the simplest way, sorting is done as before on the binary connectivity matrix and the resulting index list is applied to permutate the original connectivity matrix which holds the synaptic weights at each entry. Alternatively, the synaptic weights of the original connectivity matrix can enter the cost function by giving stronger connections more weight, this way affecting the sorting outcome itself. Future work will show to what extent our approach is generalizable to other problems in the field of matrix reordering as well.

## Material and Methods

### 1. Formulas of cost functions and their gradients

We define Δ*z*_*n,m*_ as: Δ*z*_*n,m*_ = *z*_*n*_ – *z*_*m*_ + 1

The Heaviside function H(x) as: 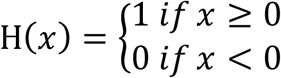

The logistic function α(*x, N*) as: 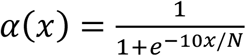

To minimize recurrent connections, we define the length term *L*(*z*) as:

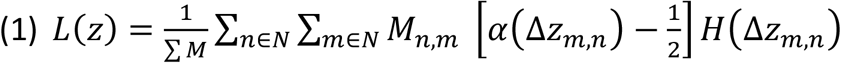

Using α^′^(*x*) = α(*x*)1 − α(*x*), the Jacobian of *L*(*z*) becomes:

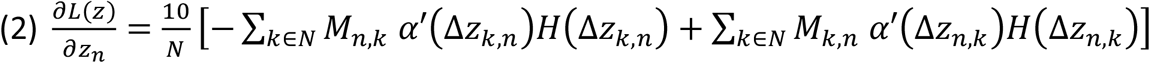

To minimize the bandwidth, we define the length term *L*(*z*) as:

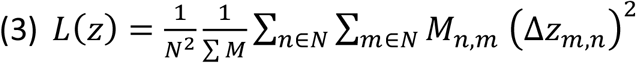

The Jacobian of *L*(*z*) becomes:

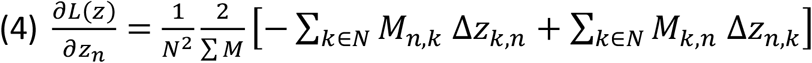

In addition, we define the Pauli term *P*(*z*) as:

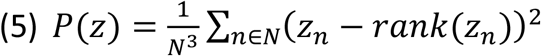

The Jacobian of *P*(*z*) becomes:

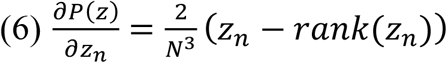

The total cost function *C*(*z*) is defined as: *C*(*z*) = *L*(z) + *P*(*z*)

### 2. Matrix generation

Original matrices, i.e. before scrambling, for the data shown in Figure 2 were constructed in the following way: elements in the lower triangle were set to 1 with a probability = 0.5, in the upper triangle with a probability = 0.04, elements along the first subdiagonal with a probability = 1. For the data in Figure 3, stripe and block matrices were constructed such that each element within the stripe or block was set to 1 with a probability = 0.5. The stripe width was set to be 40%, the block size was set to be 25% of the matrix size N.

### 3. Summary description of the smooth-index algorithm

We first turn each vertex index from a discrete integer number into a smooth real number z_i_ indicating the respective vertex position on a one-dimensional axis. Next, each vertex position is considered to be a parameter value along an independent axis. Thus, the cost becomes a function of an N-dimensional parameter space. This transition brings the advantage that the cost function can become differentiable, allowing gradient descent methods to be applied to search for a minimum. We applied this idea to reorder matrices such as a) to isolate recurrent connections, and b) to minimize the bandwidth.

#### a) Recurrency minimization

A recurrent connection is characterized by the fact that the position z_m_ of a presynaptic neuron m is larger than the position z_n_ of a postsynaptic neuron n. However, the number of recurrent connections varies in a discrete step at the point where z_m_ = z_n_, and, hence, is not a differentiable function of z. We therefore chose the average length z_m_ – z_n_ of all recurrent connections as a proxy for the number of recurrent connections. Taking a saturating function of the length reduced the stronger influence of longer connections over shorter ones. The cost function and its gradient are formulated by Eq.1 and Eq.2, respectively.

#### b) Bandwidth minimization

Here, the cost is defined as the mean of the squared length of all connections, irrespective of their sign. The respective cost function and its gradient are formulated by Eq.3 and Eq.4, respectively.

#### c) The ‘Pauli term’

To avoid all positions to collapse into a single value, we define an additional ‘Pauli term’ as the mean of the squared differences between the positions and their rank within the sorted position array. This part of the cost function and its gradient are formulated by Eq.5 and Eq.6, respectively.

The total cost is defined as the sum of the specific cost function and the ‘Pauli term’.

Both the total cost function as well as its gradient (Eqs.1,2,5,6 for minimizing recurrent connections, Eqs.3,4,5,6 for minimizing bandwidth) are passed onto the Python Scipy minimize function [29] using the bound constrained method of Broyden, Fletcher, Goldfarb, and Shanno (‘L-BFGS-B’). To return to discrete vertex indices, as the final step, the permutation list π that reorders the matrix is obtained as the vector containing the arguments of the rank-sorted parameters at the minimum of the cost function.

## Acknowledgements

We are grateful to Christian Leibold for numerous discussions. He and Juergen Haag made valuable suggestions on the manuscript.

## Notes

### Competing Interest Statement

The authors have declared no competing interest.

